# Injectable Electrospun Hydrogel with Antimicrobial, pH Sensing Nanoparticles for Local Infection Control and Monitoring

**DOI:** 10.64898/2026.07.06.736887

**Authors:** Adam Truskewycz, Shadi Houshyar, Line Pedersen, Jonathan Campbell, Braira Wahid, Jianhua Han, Ivan Cole, Peter Speck, Melanie MacGregor, Nils Halberg

## Abstract

Most antimicrobial drug candidates currently in development are derivatives of established antibiotic classes. In contrast, antimicrobial heteroatom-doped carbon quantum dot (CQD) nanoparticles vastly differ from their chemical antibiotic counterparts and exhibit potent antibacterial activity and favourable biocompatibility, representing a promising alternative strategy, particularly for topical applications. Here, we report the incorporation of cobalt-doped carbon quantum dots (Co-CQDs) into injectable, biocompatible hydrogels capable of both sensing pH and eliminating bacteria. Ultrasmall Co-CQDs demonstrated broad-spectrum activity against gram-positive Methicillin-resistant *Staphylococcus aureus* (MRSA) and Gram-negative *Pseudomonas aeruginosa* (PAO1), mediated by membrane hyperpolarisation and reactive oxygen species (ROS) induced membrane damage. The particles showed negligible effect on primary fibroblast and endothelial cell viability at concentrations that were bactericidal to MRSA. Polymeric hydrogels were fabricated via electrospinning of chitosan, polyvinylpyrrolidone (PVP), and polyvinyl alcohol (PVA) polymer blends incorporating Co-CQD and pH-responsive HPTS particles. This approach provided accurate measurement of environmental pH within the physiological range observed across healthy and chronic wounds. *In vivo*, the injectable hydrogels exhibited robust antimicrobial efficacy against MRSA without impairing wound closure relative to untreated controls, while also reducing inflammatory immune responses in infected tissues. Collectively, these findings demonstrate the potential of ultrasmall metal-doped CQDs for infection control and their integration into 3D matrices as multifunctional theragnostic platforms.

**ToC (<60 words describe the main results):** Ultrasmall antimicrobial carbon nanoparticles were incorporated into polymeric hydrogels containing a pH-responsive probe. This platform enabled detection across a physiologically relevant pH range of 5.0–6.5, spanning conditions associated with both healthy healing and chronically infected wounds. The hydrogel demonstrated strong antibacterial activity by generating damaging reactive oxygen species, and, in mice, effectively controlled infection while reducing pro-inflammatory immune responses.

Scheme 1:
Antimicrobial nanoparticle-doped hydrogel glows in alkaline wound conditions which are representative of chronic infections. Once in the wound bed, the hydrogel removes all MRSA infection, reducing inflammatory macrophage (iNOS) and neutrophil (MPO) populations, and restores healthy wound collagen deposition.

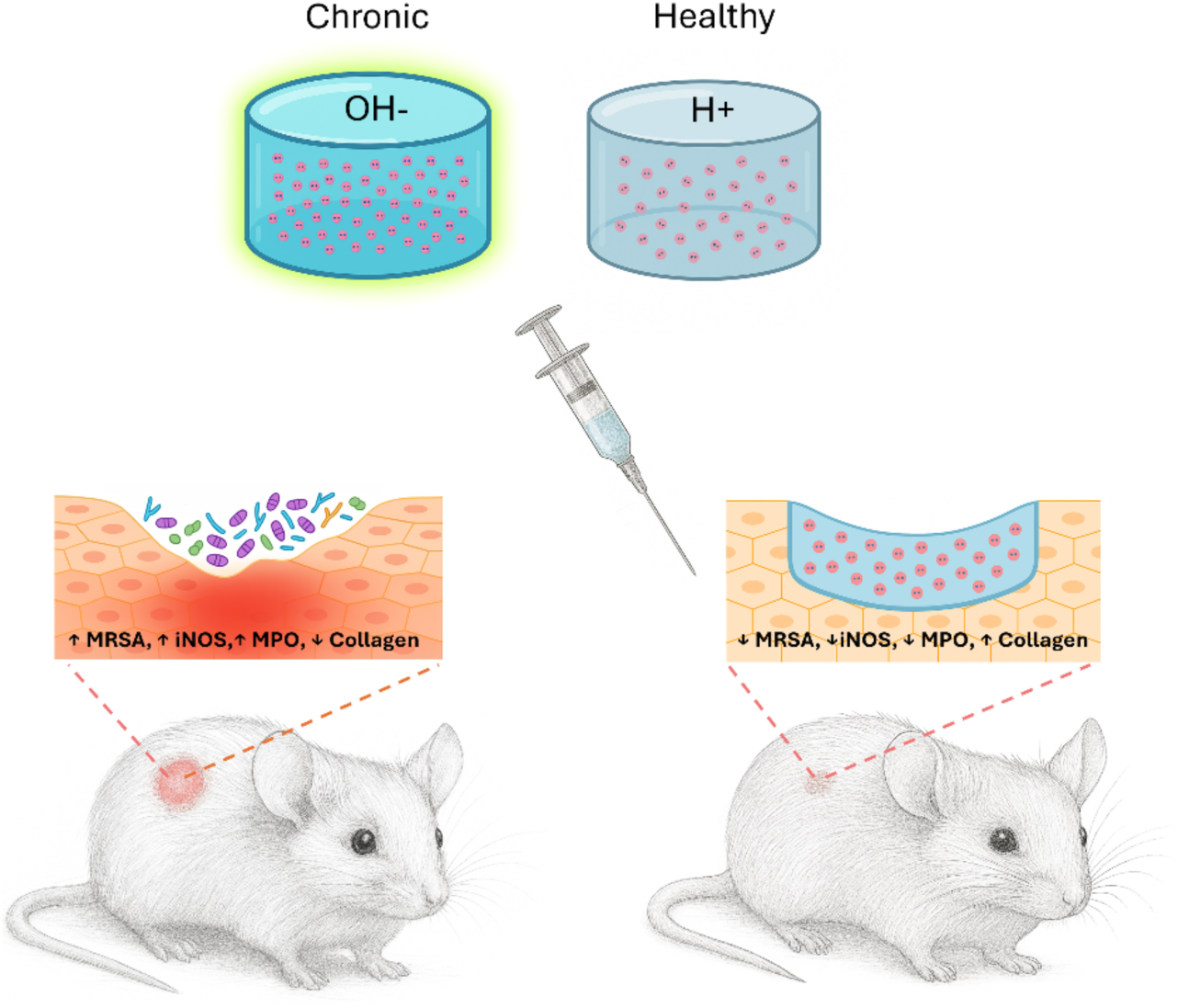

## Introduction

Bacterial infections are a major and growing global health concern, causing millions of deaths each year and placing a heavy burden on healthcare systems. In 2021, antimicrobial infections were responsible for an estimated 4.71 million deaths worldwide, while deaths involving sepsis exceeded 21 million^1^. Additionally, there is a considerable disability-adjusted life years burden among individuals living with severe microbial infections.

This challenge is further compounded by the rapid emergence of antimicrobial resistance, largely attributed to the overuse and misuse of antibiotics, together with the limited development of novel antimicrobial classes that are structurally distinct from existing drugs ^2^. Multidrug-resistant pathogens, particularly those in the ESKAPE group ^3^, including strains such as MRSA and PAO1, are increasingly responsible for difficult-to-treat hospital and community infections. Staphylococcus aureus is most frequently isolated pathogen in chronic wounds and often includes antibiotic-resistant strains like MRSA. Pseudomonas aeruginosa is a highly virulent, Gram-negative bacterium which often co-infects wounds alongside *S. aureus*, increasing the severity of the infection^4^.

Infected and chronic wounds are becoming more prevalent due to aging populations and conditions like diabetes, affecting millions worldwide and incurring substantial healthcare costs ^5^. Both acute and chronic infections lead to severe complications such as sepsis and non-healing wounds, which can result in limb amputation and increase mortality ^6,7^. Together, these challenges highlight an urgent need in wound care for alternative antimicrobial strategies beyond conventional antibiotics.

Normal wound healing progresses through four stages: clotting, inflammation, proliferation, and remodelling ^8^. During these phases, pH levels fluctuate slightly, with mildly acidic conditions (pH 5.0–6.5) helping to limit bacterial growth and support vascular regeneration, and near-neutral to slightly alkaline conditions (pH 7.0–7.5) promoting cell proliferation and tissue remodelling ^6^. In contrast, in chronic or slow-healing wounds, this process can become stalled, creating a reactive environment that impairs healing and encourages bacterial proliferation. Established bacterial populations in a wound often create an increased pH (pH 7.5->9.0) resulting from the production of ammonia, the metabolism of proteins and amino acids which release alkaline byproducts into the surrounding tissue and through the generation of alkaline proteases ^10^. The early detection of persistent infection and the tracking of their status is therefore beneficial for early intervention and to reduce complications. To address this, the integration of sensors into wound care products is being explored to monitor key biomarkers of wound health, including pH, temperature, pressure, humidity, glucose, and urea ^11,12^. Among these, pH is considered one of the most reliable indicators ^13^.

Hydrogels are widely used for dry or lightly exuding, painful, and granulating wounds, as they maintain a moist healing environment, promote autolytic debridement, and serve as a scaffold for the delivery of wound care agents ^14^. Their ability to incorporate multiple functionalities makes them attractive for a wide range of applications, including environmental sensing (e.g., pH, oxygen, and temperature monitoring) and the delivery of antimicrobial, anti-inflammatory, and pro-regenerative agents. However, it is important to determine if the electrospinning process and the subsequent incorporation of these additives within the polymeric network adversely affects their activity^15^. Unlike bandages, hydrogels, particularly injectable hydrogels, can contour to the wound’s topography, ensuring that all regions of infection are in contact with the wound care treatment ^16^. However, few injectable wound matrices combine non-antibiotic antimicrobial nanoparticles with an embedded pH readout while retaining activity after electrospinning and hydration

In addition to wound sensing, integrating antimicrobials into 3D matrices such as hydrogels can aid in fighting infections without impacting ordinary wound healing. Ultrasmall nanoparticles present a promising substitute for conventional antibiotics, especially in topical treatments where concerns about systemic bioaccumulation are minimized. Global research indicates that these nanomaterials act as versatile platforms for multifunctional activity, significantly reducing the likelihood of microorganisms developing resistance ^17^. Additionally, they can be more stably incorporated into 3D matrices than their chemical counterparts, enabling a broader range of applications.

Carbon quantum dots are ultrasmall nanoparticles known for their high biocompatibility and versatility^18^. They can be synthesised from a wide range of carbon precursors, allowing their properties to be tailored, and can also be modified to include metal dopants ^16,20^. In this study, we demonstrated that Co-CQD nanoparticles exhibit strong antimicrobial activity against both Gram-positive (MRSA) and Gram-negative (PAO1) bacteria, mediated by membrane hyperpolarisation and the generation of reactive species, including peroxidase-like activity and singlet oxygen. In parallel, a pH-responsive fluorescent probe was incorporated into chitosan nanoparticles. Both functional nanoparticle species were subsequently embedded within electrospun polymeric hydrogels composed of biocompatible polymers; Polyvinyl Alcohol (PVA), Polyvinylpyrrolidone (PVP) and chitosan, forming an injectable system capable of both monitoring wound pH and providing strong antimicrobial activity. *In vivo* studies further showed that the hydrogel platform did not impair wound healing in murine models compared to untreated controls, while effectively eliminating MRSA infections and reducing inflammatory immune responses.

## Results & Discussion

### Cobalt-Doped Carbon Quantum Dots Exhibit Antimicrobial Activity

The Co-CQD nanoparticles were synthesised using a hydrothermal method as outlined in our previous work ^21^. The resulting particles were monodispersed, with an average diameter of 2.6 nm (Figure 1A) and a slight negative surface charge of −5.4 mV (S. Figure 1A). They exhibited a dominant absorbance peak at 325 nm and a maximum fluorescence emission at 390 nm when excited at 300 nm (S. Figure 1B). X-ray photoelectron spectroscopy elemental analysis indicated that the nanoparticles were composed of carbon (65.0 At. %), oxygen (28.5 At. %), nitrogen (3.5 At. %), cobalt (1.8 At. %), and other trace elements (1.17 At. %; Figure 1B, S. Figure 1C). The concentration of cobalt was also determined to be 9.7 wt%, determined through ICP-MS analysis (Figure 1C). The presence of carboxylic acid functional groups was identified by FTIR with 3600–2650 cm^-1^ (O–H stretch), 1700 cm^-1^ (C═O stretch), 1400 cm^-1^ (O–H bend), and 1170 cm^-1^ (C–O stretch) (S. Figure 1D) and the integration of nitrogen into the matric was confirmed through XPS elemental analysis (Figure 1B, S. Figure 1C).

**Figure 1.**
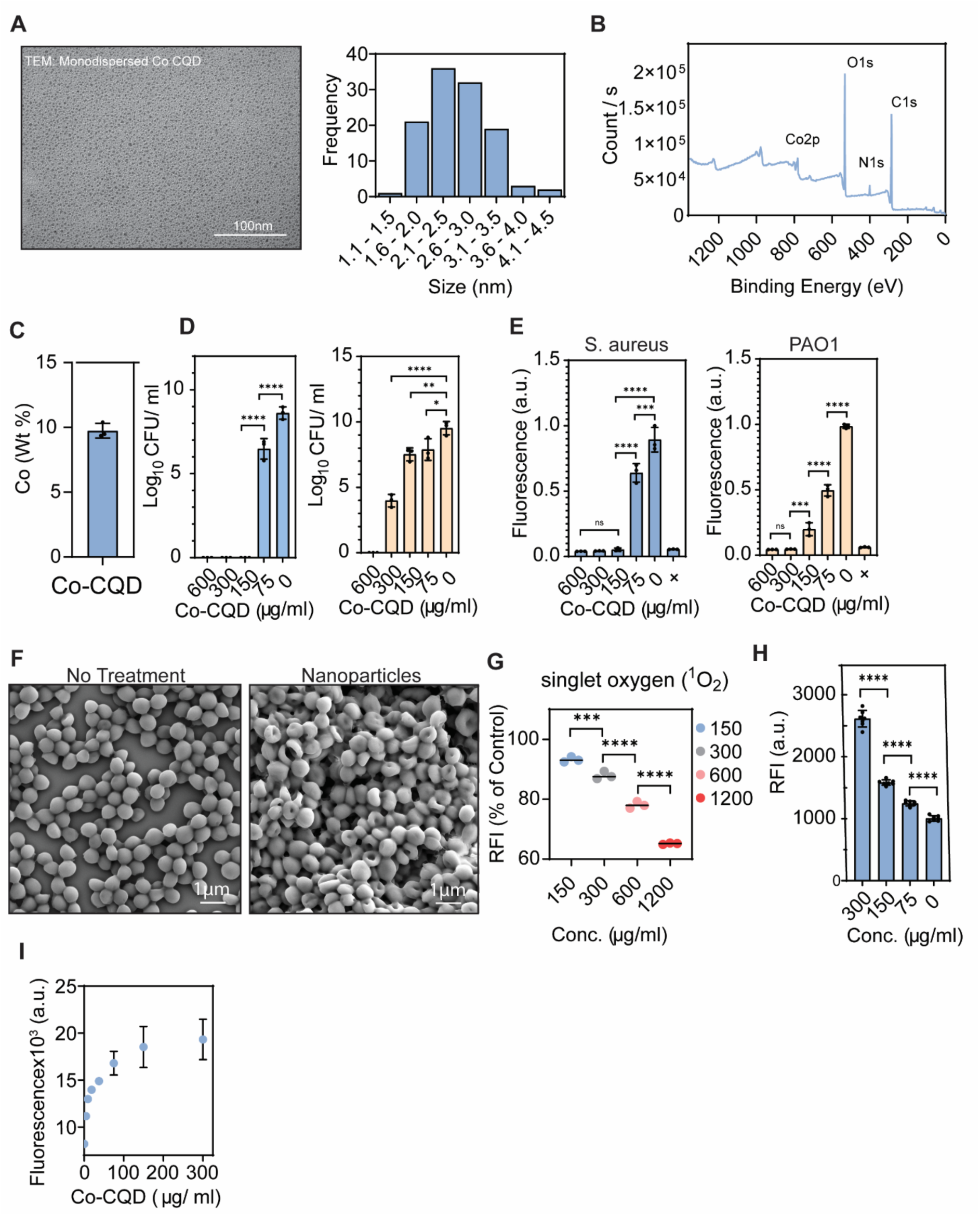
Co-CQD characterization and antimicrobial capacity. (A) Representative TEM images of Co-CQD nanoparticles (scale bar is 100 nm) and histogram showing Co-CQD size distribution. (B) X-ray photoelectron spectroscopy survey spectrum. (C) Concentration of cobalt in Co-CQD nanoparticles (Wt.%). (D) Minimum bactericidal concentrations (MBC) of MRSA and PAO1exposed to Co-CQDs. (E) Metabolic activity of MRSA and PAO1exposed to Co-CQDs. (F) Scanning electron micrograph image of S. aureus without and with exposure to Co-CQDs (scale bar 1µm). (G) Singlet oxygen production of Co-CQDs between 150-1200 µg/ml concentration). (H) Singlet oxygen production of Co-CQDs measured through the SOSG assay), (I) Peroxidase activity of Co-CQDs determined through the AMPLEX assay. Error bars represent the standard deviation. Statistical significance was determined using the one-way ANOVA with Tukey’s multiple comparison test. ns, *, **, ***, **** signifies not significant, p < 0.05, p < 0.005, p < 0.0005 and p < 0.0001, respectively.

The Co-CQD particles exhibited potent antimicrobial activity against both *S. aureus* and PAO1, with minimum bactericidal concentrations of 150 µg/ml and 600 µg/ml, respectively (Figure 1D). This activity was further supported by a halting of metabolic activity in both bacterial species at nanoparticle concentrations of 150 µg/ml for *MRSA* and 300 µg/ml for PAO1, with noticeable reductions beginning at 75 µg/ml and higher for both species (Figure 1E). The antimicrobial effect was more pronounced in the gram-positive MRSA, prompting additional mechanistic investigations using this organism. This was reinforced through the particle’s increased antimicrobial activity against *Enterococcus faecalis* over *Escherichia coli* (S. Figures E, F, G, and H).

### Cobalt-doped Carbon Quantum Dots Hyperpolarise Bacterial Membranes and Generate Damaging ROS

Scanning electron microscopy shows that *MRSA* subjected to nanoparticles possess a cavitated morphology, indicating structural membrane deformations (Figure 1F). Bacterial membrane damage was supported by increased extracellular alkaline phosphatase release from cells exposed to Co-CQD nanoparticle concentrations above 30 µg/ml (S. Figure 1I), and membranes were increasingly hyperpolarised with increasing Co-CQD concentration (S. Figure 1J). Transmission electron microscopy also shows cellular debris resulting from Co-CQD treatment (S. Figure 1K).

The particles showed capacity to generate both singlet oxygen (^1^O_2_) through both ABDA and SOSG assays (Figure 1G, H). The ABDA assay showed significant concentration-dependant ^1^O_2_ production between 150-1200 µg/ml, and the SOSG assay showed increased ^1^O_2_ production between 75-300 µg/ml. In addition to singlet oxygen, the particles were shown to generate ROS, through the breakdown of peroxides (peroxidase assay) and oxidise ascorbic acid at low particle concentrations (12.5-50 µg/ml; Figure 1I, S. Figure 1L).

These findings are in good agreement with our previous work^21^ where the antimicrobial activity of cobalt doped carbon quantum dots against *Escherichia coli*, *Enterococcus faecalis* and *Staphylococcus aureus* was synergistically enhanced in a weak acetic acid environment.

### HPTS-Chitosan pH-Sensing Nanoparticle Formation

8-Hydroxypyrene-1,3,6-trisulfonic acid trisodium salt (HPTS) dye is a biocompatible, water-soluble, and highly sensitive pH probe; however, because it lacks suitable functional groups for stable covalent attachment, its integration into 3D matrices is limited ^22^. In this study we bound HPTS to chitosan and transformed this complex into stably fluorescent pH-responsive nanoparticles.

Chitosan was first dissolved in acetic acid (0.75 M), after which the pH-responsive HPTS was incorporated. The strong electrostatic interaction between the sulfonate groups of HPTS and the protonated amino groups of chitosan has been previously reported in the literature and shown to facilitates electrostatic complex formation ^23^. The chitosan-HPTS mixture in acetic acid was then subjected to hydrothermal treatment to promote grafting/polymerisation between the interacting species and enhance the hydrophilicity of the resulting complex. Due to HPTSs high thermal stability ^24^, hydrothermal conditions did not deactivate its fluorescent and pH sensing properties. The pH-responsive nanoparticles generated during this process (Figure 2A) were then filtered and purified by dialysis. These particles were incorporated into electrospun polymeric gels, with and without antimicrobial Co-CQD nanoparticles.

**Figure 2:**
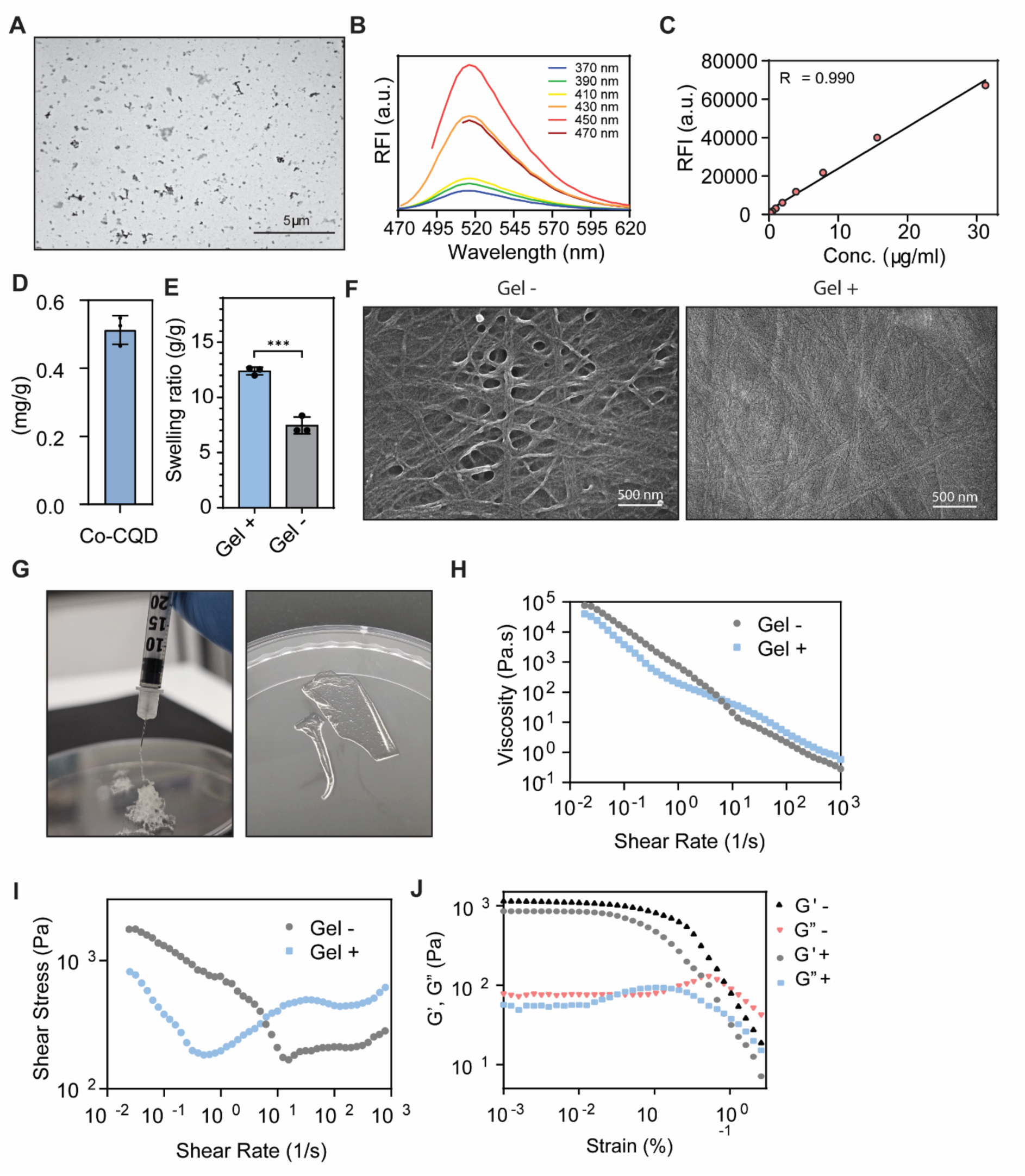
Fluorescent HPTS nanoparticle and hydrogel characterisation. (A) Representative transmission electron microscope image of HPTS/chitosan nanoparticles. (B) HPTS/chitosan fluorescence excitation and emissions profile. (C) Linear region of HPTS/chitosan nanoparticle fluorescence relative to particle concentration. (D) Concentration of Co-CQDs incorporated into hydrogel. (E) Water absorbing capacity of freeze-dried hydrogels with and without Co-CQDs. (F) SEM images of hydrogel without and with Co-CQDs. (G) Photographs of Co-CQD containing hydrogel being passed through a needle and in a thin film. (H) Shear thinning property of hydrogel with and without Co-CQDs using the flow step method. (I) Stress ramp to detect onset of material/hydrogel flow using the flow step method. (J) Oscillatory stress sweeps of hydrogel in solid state. Error bars represent the standard deviation. +, - Gel refers to the hydrogel containing Co-CQD nanoparticles (+) and hydrogel without Co-CQD nanoparticles (-) respectively. Statistical analysis of hydrogel swelling capacity was determined via an unpaired t-test with Welch correction. ns, *, **, *** signifies not significant, p < 0.05, p < 0.005, p < 0.0005 respectively.

The HPTS/chitosan nanoparticles possessed a main absorption peak at 450 nm with a linear absorbance response to pH between 0.25 – 31.25 µg/ml (S. Figure 2A) and had an optimum fluorescence excitation wavelength at 450 nm resulting in a fluorescence emissions peak of 515 nm (Figure 2B). The composite showed concentration-dependent absorbance and fluorescence output at physiological pH (pH 7.4) and linear absorbance and fluorescence profiles were observed between 0-125 µg/ml (R^2^= 0.666) and 0- 30 µg/ml (R^2^= 0.660; Figure 1C) respectively. From this data, a composite concentration within the absorbance and fluorescence linear region was chosen to incorporate into electrospun polymeric gel fibres.

The FTIR spectrum of the HPTS/chitosan nanoparticles (S. Figure 2B) did not display any clear characteristic peaks attributable to HPTS, particularly the intense band in the 1165–1140 cm⁻¹ range associated with asymmetric S=O stretching of sulfonate (–SO₃⁻) groups, suggesting that these groups are not detectable within the particles. Nevertheless, following extensive purification consisting of repeated heating to 70 °C, centrifugation, and subsequent dialysis, the nanoparticles still exhibited fluorescence characteristic of the dye without the presence of any fluorescent component from unbound dye leaching into the supernatant.

### Electrospun Hydrogel Fabrication and Mechanical Properties

Antimicrobial, pH-responsive electrospun polymeric fibre/gels were produced by electrospinning a blend of chitosan, PVP, and PVA at a 1:2:2 w/w ratio, incorporating Co-CQD nanoparticles (5 mg/g) and the chitosan–HPTS nanoparticles (5 µg/ml) (Figure 2D). A solution of methanol, acetic acid and water was required to solubilise all polymers and facilitate the incorporation of nanoparticles for the electrospinning process. The freeze-dried hydrogels (S. Figure 2C) incorporating Co-CQD nanoparticles absorbed and retained 12.4 times their dry weight in water after 24 h of hydration, whereas hydrogels without Co-CQD additives exhibited a markedly lower water-holding capacity of 8.3 times their dry weight (Figure 2E). An increased cross-link density generally reduces swelling while enhancing mechanical strength. These results suggest that the incorporation of Co-CQD nanoparticles disrupts the repeated polymer cross-linking present, reducing network strength, but enhancing water uptake and retention. Following synthesis, matrices containing Co-CQDs showed a less obvious fibrous morphology than gels without Co-CQD incorporations (Figure 2F).

The hydrogel’s mechanical behaviour in its Injected and solid state (Figure 2G) was characterised. The viscosity of the material was measured using increasing shear rates and the viscoelastic properties (G’, G”) were measured at 1Hz as a function of applied strain (S. Figure 2D). The addition of Co-CQD nanoparticles to the gel resulted in a weaker injectable gel as shown by lower viscosity (Figure 2H) and shear stress at shear rates < 6s^-1^ (Figure 2I). However, at higher shear rates (above 6s^-1^) the trend is reversed, where the injectable gel containing Co-CQD nanoparticles has higher viscosity (and shear stress) in the flow experiment. This may be due to some reorganisation or formation of new interactions within the gel caused by the Co-CQD nanoparticles functional groups, or as the hydrogel becomes more fluid under increasing shear (shear thinning), the nanoparticles flow relatively less freely and become “jammed” by their neighbours. This aggregation results in an increase in the Co-CQD containing gel’s relative viscosity^25^.

Figure 2J display the elastic (G′) and viscous (G″) shear modulus curves obtained from strain amplitude sweeps for the control hydrogel and the Co-CQD–containing hydrogel not passed through a syringe. At low strains below 0.1%, the control gel maintains its structural integrity, as indicated by G′ and G″ remaining within the linear viscoelastic region. In this system, G′ exceeds G″, confirming that the gel exhibits predominantly solid-like behaviour. When strain increases beyond the range of 0.01-0.1%, the gel structure begins to break down, evidenced by a decrease in G′ and the appearance of a peak in G″. At approximately 1% strain, the yield point is reached, indicated by the crossover of G′ and G″.

The Co-CQD–containing hydrogel exhibiting a weaker structural network than that of the control with deformations starting to occur at a lower strain values compared to the gel without Co-CQDs. This early onset of breakdown was reflected by a reduction in G′ and the emergence of a peak in G″. The G” peak was broader than that of the control indicating that the loss of the elastic structure occurs over a wider range of deformations. This could be due to a more complex network structure in this sample, and is commonly observed in soft gels where an interconnected network of cross-linked particles exists^26^. A weaker structural network improves hydrogel injectability (Figure 2G), which is advantageous for filling and conforming to irregularly shaped infected wound beds.

### pH Responsive Behaviour of Antimicrobial Hydrogel in Wound Relevant Media

The electrospun gel was evaluated for its pH-dependent absorbance and fluorescence behaviours. Absorbance measured at 450 nm increased steadily from pH 5 to pH 9, demonstrating a clear and distinguishable pH response across the mildly acidic to mildly alkaline range (Figure 3A, B). The polymer also exhibited a highly sensitive fluorescence response to pH (Figure 3C), with resolvable shifts occurring in 0.5 pH increments between pH 5 and 9.5 (Figure 3D, E), covering the full pH spectrum typically observed in healthy versus chronic wound environments with the latter frequently possessing highly alkaline pH resulting from by-products of bacterial metabolism (i.e. production of ammonia, metabolism of proteins and amino acids and the generation of alkaline proteases etc.). The pH fluorescence intensity has a sigmoidal relationship with R^2^=0.667 representing an extremely strong S-shaped (logistic) data fit (Figure 3E).

**Figure 3:**
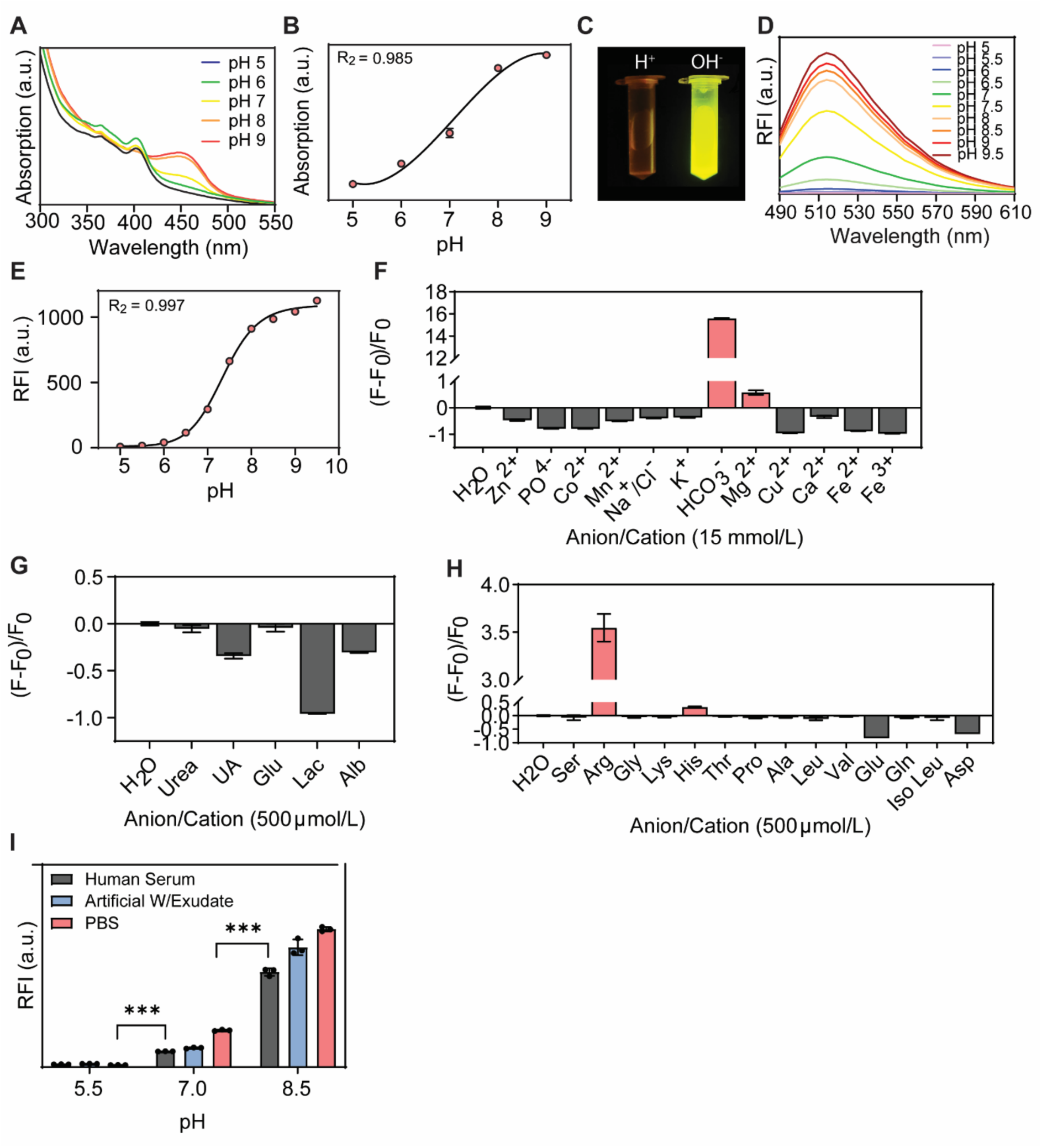
Hydrogels pH sensing properties. (A) Hydrogels absorbance spectra within the pH range of 5-6. (B) Endpoint absorption intensity of hydrogel within the pH range of 5-6. (C) photograph of hydrogels fluorescent component in acidic (pH 5.5) and basic (pH 8.5) conditions. (D) Fluorescence spectra of hydrogel in differing environmental pHs between pH 5.0-6.5. (E) Relationship between pH and fluorescence endpoints of hydrogel at 514 nm wavelength. (F) Fluorescence intensity of hydrogel in the presence of varying anion/cations found in chronic wounds. (G) Fluorescence intensity of hydrogel in the presence of varying chemical entities found in chronic wounds. (H) Fluorescence intensity of hydrogel in the presence of various amino acids found in chronic wounds. (I) Influence of human serum, artificial wound exudate, and phosphate buffered saline (PBS) on hydrogels fluorescence intensity. Error bars represent the standard deviation. Statistical significance was determined using the one-way ANOVA with Tukey’s multiple comparison test. ns, *, **, *** signifies not significant, p < 0.05, p < 0.005, p < 0.0005 respectively.

Resolving pH in an interference-free environment does not accurately reflect the conditions present in a chronic wound. To address this limitation, the gel was challenged with a range of anions and cations commonly found in chronic wound exudate ^27^. Of the twelve ionic species assessed (Figure 3F), only two caused an increase in fluorescence consistent with a shift toward alkaline conditions. Magnesium which is known to support cell migration, collagen synthesis, and angiogenesis produced minimal interference with an increase in fluorescent signal with a (F–F₀)/F₀ value of <1. In contrast, bicarbonate (HCO₃⁻) induced a substantial fluorescence increase, with a (F–F₀)/F₀ value >15. Bicarbonate acts as a physiological buffer that supports blood flow, oxygenation, and cell viability within the wound environment. It is important to note, however, that in situ this ion is not present in isolation; interactions among multiple ionic species collectively influence wound pH.

Several organic species found in slow healing wounds such as urea, uric acid, glucose, lactate and albumin were assessed for activity with pH probe with all species showing a decreased fluorescence of the polymeric matrix (Figure 3G). Additionally, the impact of 14 different amino acids was assessed on the gel’s fluorescence sensing capabilities (Figure 3H). Of these only histidine and arginine gave an increase in fluorescence. The significant increase in fluorescence from arginine ((F–F₀)/F₀ of ∼3.5) can be attributed to its strongly basic nature. Arginine is responsible for collagen deposition, increased blood flow and stimulating growth hormones in a wound environment. Histidine is considered to be a basic amino acid but acts as a buffer in biological systems. Its presence led to a minor increase in fluorescence ((F–F₀)/F₀ of <0.5).

After evaluating each potential interfering species independently, it became clear that those producing increases in fluorescence were all basic in nature. Consequently, in a physiologically relevant mixture, the presence of additional ions with differing pH influences would likely mask the impact of these alkaline species on the sensor’s signal. In practical terms, the pH sensor reflects the overall environmental pH; therefore, to investigate this further, the polymer’s fluorescence responsiveness was examined at pH 5.5, 7.0, and 8.5 in three different environments: PBS, artificial wound exudate (containing a mixture of anions, cations, amino acids, and other species in concentrations typical of chronic wound fluid), and human serum. Under neutral and alkaline conditions, variations in fluorescence intensity arose solely from differences in medium composition, with PBS producing the highest signal, followed by artificial wound exudate, and then human serum. Importantly, despite these baseline differences, the fluorescence profiles for each pH remained distinct with no overlap between media type. This demonstrates that the polymeric gel can clearly discriminate between acidic, neutral, and alkaline conditions regardless of the wound-representative environment (Figure 3I).

### Electrospun Hydrogels Maintain Bactericidal Activity *in vitro* and *in vivo*

Polymeric gels with and without Co-CQDs were assessed for their antimicrobial activity through a live/dead assay where the compromised cell membranes of dead cells uptake membrane-impermeant propidium iodide dye, staining them with the red fluorescing dye. Gels containing the nanoparticles caused 94.4 % MRSA death when incubated for 24 h where those without nanoparticles had >93 % viability (Figure 4A). MRSA morphology on fibres containing nanoparticles had a shrivelled and bumpy morphology whilst those on fibres without nanoparticles were smooth and spherical (Figure 4B). This reinforces the Co-CQDs damaging effect of on bacterial membranes.

**Figure 4:**
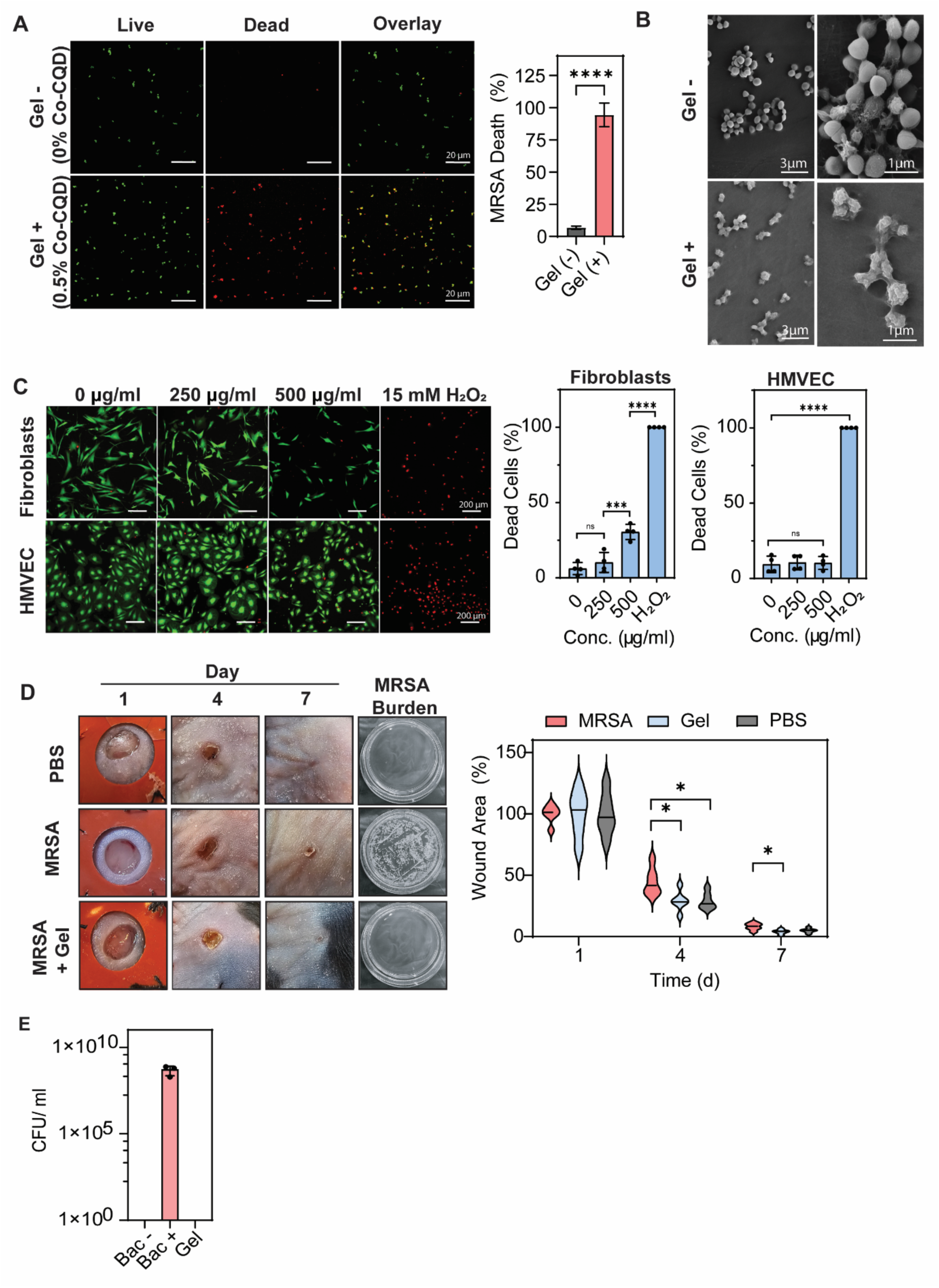
Antimicrobial Activity and Biocompatibility of antimicrobial hydrogel. (A) Representative fluorescent confocal microscope images of MRSA in the presence of differing Co-CQDs concentrations. The live/dead probes fluoresce green in viable cells and red in dead cells. (B) SEM images of MRSA on hydrogel without and with Co-CQDs. (C) Representative fluorescent confocal microscope images of human dermal fibroblasts and human dermal microvascular endothelial cells in the presence of differing Co-CQDs concentrations. The live/dead probes fluoresce green in viable cells and red in dead cells. Quantification of live and dead cells accompanies these images. (D) Representative images and graphical representation of full thickness wounds in Black C57BL/6 mice, treated with PBS, infected with MRSA and then treated with hydrogels for 7 d, along with (E) bacterial burden of wounds after 7 d.

The Co-CQD containing gel was added to dermal fibroblast and dermal microvascular endothelial cells with Co-CQD concentrations at 500 µg/ml and 250 µg/ml. At 500 µg/ml Co-CQD concentration, there was approximately 30 % cell death after 24 h which dropped to below 10 % at 250 µg/ml concentration. Surviving fibroblast cells that were exposed to high concentrations of Co-CQDs were able to recover when their growth medium was replaced with fresh, unamended media (S. Figure 3A). Human microvascular cells were more resilient to Co-CQD doped gels with both 500 and 250 µg/ml Co-CQD concentrations resulting in less than 10% cell death (Figure 4C).

Having demonstrated that the Co-CQDs could kill bacteria in a free and immobilised state, their effectiveness was assessed *in vivo*. Wounds were made onto the back of female Black C57BL/6 mice, splinted and MRSA bacteria was inoculated at 2.7 × 10^6^ CFU. The wounds were then allowed to dry for up to 3 hours before being treated with electrospun gels containing CQDs (100µl of gel). A PBS control without bacteria or gel was used to assess an uninfected wound healing environment. After 7 days there was no evidence of bacterial infection from the gel containing CQDs (Figure 4D, E). Mice wounds infected with MRSA but without gel treatment showed strong bacterial proliferation containing bacterial concentrations above 10^8^ CFU/ml after the 7-day exposure period.

Over 7 days, mice weights remained steady regardless of treatment indicating that the treatment was well tolerated (S. Figure 3B). Wound closure rates between CQD gel and the PBS control showed a slightly elevated wound closure rate on days 4 and 7 indicating that the gel and CQD containing gels was well tolerated and promoted normal wound healing.

### Co-CQD Containing Hydrogels Normalize Infected Wound Immune Environment

To support these observations, immunofluorescence staining was performed on wound tissue after 7 days of healing to assess the presence of macrophages (F4/80), inflammatory macrophages (iNOS), and neutrophils (MPO) (Figure 5A, B). In MRSA-infected wounds treated with Co-CQD gels, a significant decrease in total macrophage populations were observed compared the MRSA control, as assessed by F4/80 expression (Figure 5A, B). There was also a significant difference in the proportion of pro-inflammatory macrophages, as evidenced by increased iNOS expression. This suggests that the hydrogel effectively eliminated the MRSA responsible for inducing the pro-inflammatory M1 macrophage phenotype, while not triggering an inflammatory response on its own which is beneficial for resolving inflammation, clearing cellular debris, and driving tissue repair ^28^. Uninfected wounds treated with PBS and MRSA-infected wounds treated with the Co-CQD hydrogel showed significantly reduced neutrophil concentrations compared to the MRSA control observed through reduced MPO presence (Figure 5 A, B). Lower MPO has been linked with wounds possessing decreased oxidative damage, increased collagen deposition and increased healing ^26^.

**Figure 5:**
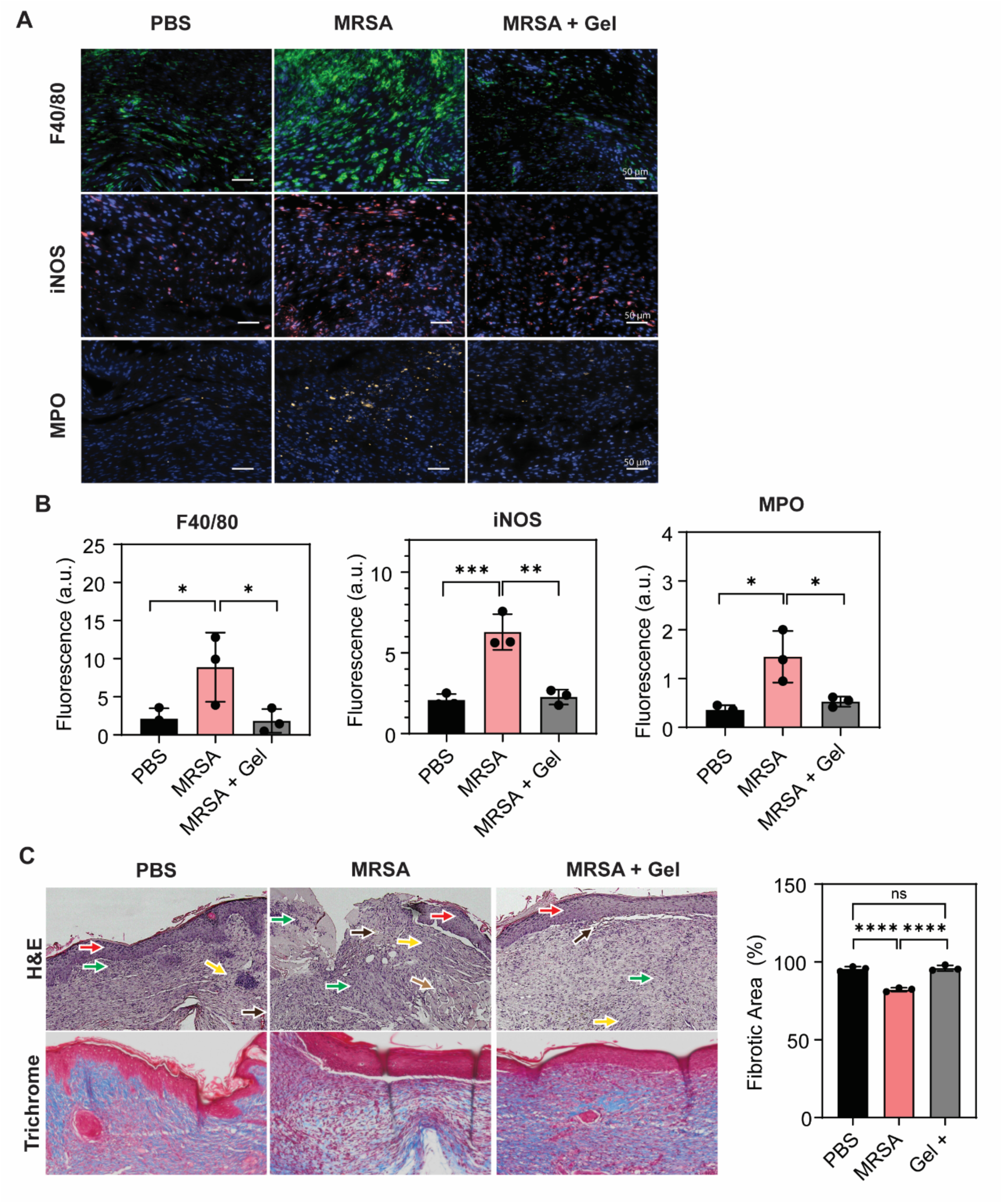
***In vivo* cell and tissue compatibility.** (A) Confocal microscopy images of fluorescent antibody labelled F4/80 macrophage, iNOS inflammatory macrophage, and MPO neutrophil presence in wound areas following treatment after 7 d. (B) Quantification of F4/80 macrophage, iNOS inflammatory macrophage, and MPO neutrophil presence in wound areas following treatment after 7 d. (C) Representative H&E-stained wound regions of PBS, MRSA-infected, and hydrogel-treated MRSA-infected wounds. Green arrows denote mixed inflammatory/immune cells, Red arrows denote the epidermal layer, Brown arrows denote haemorrhagic defects, Black arrows denote neovascularization, and yellow arrows denote fibroblast cells. Trichrome staining shows collagen deposition from treatments. Error bars represent the standard deviation. Statistical significance was determined using a one-way ANOVA with Tukey’s multiple comparison test. ns, *, **, ***, **** signifies not significant, p < 0.05, p < 0.005, p < 0.0005 and p < 0.0001, respectively.

In Hematoxylin and Eosin (H&E) sections (Figure 5C), staining of untreated infected wound areas revealed haemorrhagic defects (brown arrow), an incomplete epidermal layer (red arrow), fibroblast cells (yellow arrow), and regions of densely concentrated mixed inflammatory/immune cells (green arrows). Granulation tissue is a specialised, temporary connective tissue formed during the proliferative phase of wound healing and contains a significant population of immune cells alongside structural cells (Figure 5C). Granulation tissue was observed across all samples; however, it appeared more extensive in the untreated MRSA control.

Fibroblasts were present across all wound groups regardless of treatment, however, Masson’s trichrome staining revealed reduced collagen deposition in MRSA-infected wounds, suggesting impaired fibroblast function (Figure C). In contrast, treated groups exhibited more robust collagen formation, indicative of a shift toward a proliferative and reparative fibroblast phenotype and reduced neutrophil numbers, as observed in the decrease of MPO (Figure 5 A, B).

Wound healing involves the formation of fibrous tissue (collagen) to repair damage^30^. Healing mouse wounds subjected to gel treatments had significantly greater collagen content when compared to MRSA infected untreated wounds as indicated through binding to the blue Masson’s stain, showing similar collagen formation to the PBS control (Figure 5C). An infected wound typically shows reduced collagen formation, resulting in weaker tissue, because bacteria and extended inflammation degrade collagen faster than it can be regenerated ^31^. The gel treatment in this study effectively eliminated MRSA infection, thereby aiding in the restoration of normal collagen formation. Carbon dots, both alone and when incorporated into hydrogel matrices, have previously been demonstrated to enhance collagen deposition following the treatment of infected wounds^32,33^.

## Conclusions

In summary, a biocompatible injectable hydrogel incorporating both pH-sensitive and antimicrobial nanoparticles was successfully developed. This platform demonstrated stepwise pH sensing across conditions representative of normal healing (pH 5.0-7.5) and infected chronic wounds (pH > 8). It effectively cleared MRSA infections in a mouse wound model, reducing levels of inflammatory neutrophils and macrophages. Wounds treated with the hydrogel healed at rates comparable to uninfected controls and supported normal collagen deposition, reinforcing the materials biocompatibility and unhindered influence on healing processes. Overall, this system demonstrates strong potential as an advanced wound dressing, allowing patients to monitor the condition of slow-healing wounds through pH sensing, while also managing infection and preserving a moist environment that supports healing. Future work should address long-term stability and sensor calibration in diabetic chronic wound models.

## Methods

### Chemicals Used in this Study

8-Hydroxypyrene-1,3,6-trisulfonic acid, trisodium salt (HPTS), ZnSO_4_ · 7H_2_O, KH2PO4, CoCl_2_ 6H_2_O,MnCl₂, NaCl, KCl, NaHCO₃, MgCl_2_, Cl_2_CuH_4_O_2_, CaCl_2_-2H_2_O, FeCl₂, FeCl_3_ · 6H_2_O.

### Synthesis of HACC CQD’s

Cobalt-doped carbon quantum dots were synthesized through a hydrothermal method. In a typical synthesis, citric acid (5.4 g) and hexamine cobalt chloride (1.0 g) were dissolved in 30 ml of ultrapure water (Milli-Q) inside a Teflon-lined autoclave. The solution was heated to 200°C for 2 h and the resulting solution was purified via dialysis (100-500 MWCO) for 48 h before being lyophilised to a powder.

### Characterization of CQDs

Nanoparticles were imaged via a transmission electron microscope (Hitachi HT7800) at 100kV voltage. Particle fluorescence and absorbance was measured using a Tecan SPARK® in flat bottom 66 welled plates (Costar). X-ray Photoelectron Spectroscopy (XPS) measurements were obtained using a Thermo Scientific K-Alpha with a monochromated Al 1487 eV Kα source and sample composition was determined using CasaXPS 2.3.23 software. Particle cobalt concentration was measured from acid-digested samples in an Element XR (Thermo Scientific) ICP-MS. FTIR analysis was performed using a Bruker VERTEX 80v UHV FTIR under vacuum between 400-4000cm^-1^ wavenumber range. Nanoparticle zeta potential was measured in Milli-Q water using a Malvern 2000 Zetasizer within a DTS 1060C cuvette (Malvern).

### Synthesis of HPTS/Chitosan Nanocomposite

8-Hydroxypyrene-1,3,6-trisulfonic acid, trisodium salt (HPTS) (0.05g), chitosan (0.5 g), and acetic acid (66%) (3 ml) + H_2_O (37 ml) were transferred to poly(tetrafluoroethylene)-lined autoclaves (50 ml). Vessels were heated at 200°C for 2 h and cooled to room temperature. The resulting solution was filtered through 0.2 μm syringe filters, subjected to repeated heating (70°C) and centrifugation cycles, and subsequently purified by dialysis (100–500 MWCO) against ultrapure water for 50 h with regular water replacement. Nanoparticle stability was confirmed when no fluorescence attributable to the water-soluble parent dye molecule was detected in the centrifuged supernatant at λ_ex = 450 nm and λ_em = 515 nm. The purified solution was subjected to lyophilisation to obtain a powdered product.

### Determination of HPTS in HPTS/Chitosan Composite

HPTS/chitosan NPs and HPTS alone were made to different concentrations (µg/ml) in PBS adjusted to pH 7. Fluorescence values were obtained from λex = 450 nm and λem = 515, and absorbance values were obtained at OD_450_. Relative fluorescence intensity of HPTS/ chitosan composite/ mg compared to HPTS/ mg alone was used to determine concentration of HPTS present in composite.

### Electrospun Hydrogel Synthesis

For the base polymer formulation, chitosan, PVP, and PVA were mixed at a 1:2:2 weight ratio and dissolved in a solvent system composed of 30% aqueous acetic acid and methanol at a 1:1 volume ratio. The polymer solution was supplemented with chitosan/HPTS nanoparticles at a final concentration of 5 µg/ml, selected from the linear region of the concentration-dependent fluorescence standard curve. Co-CQD nanoparticles were then incorporated at 5 mg/g. The resulting mixture was stirred until homogeneous and subsequently sonicated for 5 minutes to improve particle dispersion. Fibrous gel matrices were fabricated by electrospinning the prepared solution through a 20G needle at an applied voltage of 13.5 kV and a flow rate of 0.05 ml/min. The tip to collector distance was maintained at 32 cm. To facilitate handling during fabrication, a polycaprolactone (PCL) support layer was prepared from 5% PCL in methanol and electrospun as a backing layer. The PCL support was removed prior to subsequent experimental use.

### Hydrogel Swelling Capacity

Freeze-dried hydrogels were initially weighed and then immersed in water for 12 hours to allow swelling. After incubation, excess surface water was carefully removed using Kimwipe tissue. The swollen hydrogels were weighed again, and the water holding capacity was calculated by comparing the initial dry weight to wet weight gain (g/g).

### Hydrogel Rheology

The rheological behaviour of the hydrogels was measured with a TA Instruments AR2000 rheometer using a flat 25 mm aluminium plate. Hydrogels with and without Co-CQD nanoparticles were prepared as disk-shaped samples with a 25 mm diameter and ∼1 mm thickness prior to testing. Measurements of the viscoelastic properties such as storage modulus (G′) and loss modulus (G″) were conducted within the linear viscoelastic region under oscillatory conditions at 1Hz from 0.02 to 3000% strain. Additional rheological tests were performed on the hydrogel once it had passed through a syringe needle. Viscosity was determined using the step-flow method using a continuously rotating plate with a gap of 250µm over a range of shear rates from 0.01 to 3000 s^-1^ on 200 µl of gel injected through a 20G needle.

### Fluorescence Measurements

The fluorescence response of different cations to the Co-CQD/HPTS-doped hydrogel was investigated to determine its selective nature. The gel (100 µl) was added to wells of black 96-well plates. Anions/ cations typically found in chronic wound environments were added to the solution at a final concentration of 15 mmol/L (Mg^2+^, Cu^2+^, Fe^3+^, Fe^2+^, Zn^2+^, K^+^, Na^+^, Mn^2+^, Ca^2+^, Co^2+^, PO_4_^3-^, HCO_3_^-^). Additionally, amino acids implicated in wound/ chronic wound settings were also added to the probe at 500 μmol/L concentration (Serine, Arginine, Glycine, Lysine, Histidine, Threonine, Proline, Alanine, Leucine, Valine, Glutamic acid, Glutamine, Iso Leucine, Aspartic acid). Further wound/ chronic wound relevant chemicals were tested for fluorescence interaction with the probe at 500 μmol/L concentration (urea, uric acid, glucose, lactate, albumin). Fluorescence measurements were performed at λex = 450 nm and λem = 515 nm. Fluorescence intensity was represented by (F-F_0_)/F_0_ where F = fluorescence intensity of the gel in the presence of the treatment and F_0_= fluorescence intensity without treatment.

Following the investigation of individual potential competing species on the gel separately, the gel was added to human serum (Sigma Aldrich) and artificial wound exudate to assess how numerous potential competing species may influence the fluorescence of the gel at differing pHs (pH 5.5, 7, and 8.5). The artificial wound exudate comprised of chemical species found in chronic wound environments (mmol/ L) ^34,35^. Consisting of Zn^2+^ (0.1), PO_4_^-^ (1.47), Co^2+^ (0.1), Mn^2+^ (0.006), Na^+^ (146), K^+^ (6.6), HCO_3_^-^ (31), Mg^2+^ (1.3), Cu^2+^ (0.03), Ca^2+^ (5), Fe^2+^ (1.3), Fe^3+^ (1.3), Serine (0.360), Arginine (0.432), Glycine (0.511), Lysine (0.607), Histidine (0.137), Threonine (0.221), Proline (0.373), Alanine (0.626), Leucine (0.403), Valine (0.361), Glutamine (0.401), glutamate (0.451), Isoleucine, (0.164), Aspartate (0.066), Urea (23.1), Uric acid (0.823), Glucose (6.6), Lactate (6.5), Albumin (50 mg/ml).

PBS served as the control. The gels interaction in varying environments was represented as relative fluorescence intensity.

### Cellular Toxicity

Human Dermal fibroblasts (dFIB) (CC-2509- LONZA) were cultured in DMEM (10% FBS and 3% glutamine) and Human Dermal Microvascular Endothelial cells (HMVEC-dAd) (CC-2543-LONZA) were cultured in EBM™-2 media supplemented with EGMTM-2 Endothelial SingleQuots^TM^ Kit. Cells were grown to above 50% confluency in IBIDI 8 welled plates and hydrogel containing Co-CQD nanoparticles were added at different nanoparticle concentrations (0, 250 & 500 µg/ ml) or positive killing control (15 mM H_2_O_2_) and incubated for 24 h at pH 7. Cytotoxicity was determined by a LIVE/DEAD® Cell Imaging assay (R37601-Invitrogen). Briefly, the kit reagents were added to cells as per the manufacturer’s suggestions, incubated for 20 minutes and imaged on a Dragonfly 505 (Andor Technologies, Inc) using 60 X objectives with oil immersion. Images were taken using excitation/ emission wavelengths 488/515 and 570/ 600 nm.

### TEM for Co-CQD’s on Bacterial Cells

Samples were fixed in 2,5% glutaraldehyde (diluted in a 0,1 M sodium cacodylate buffer) and 2% paraformaldehyde for 24 hrs at 4°C. Post fixation was performed for 1 h (on ice) in 1% osmium tetroxide (EMS # 19134) diluted in 0,1 M sodium cacodylate buffer, followed by 2 washing steps. The samples were then dehydrated using a graded ethanol series (30%, 50%, 70%, 96% and 100%) before being transferred to a 1:1 solution of 100% ethanol: propylene oxide (15 min). Samples were then transferred to 100% propylene oxide (15 min) before gradually introducing agar 100 resin (AgarScientific R1031) drop by drop over the next hours. Samples were then transferred to a small drop of 100% resin and excess propylene oxide was allowed to evaporate (1h). Samples were then transferred to 100% resin and placed in molds and left in room temperature overnight. The molds were then placed at 60℃ for 48h to polymerize. Ultrasections of approximately 60nm were placed on 100 mesh formvar coated (EMD # 15820) copper grids (EMS #G100H-Cu) and stained with 2% uranyl acetate (EMS # 22400) and lead citrate (VWR #1.07398). Grids were imaged using a Hitachi HT7800 transmission electron microscope at 100kV.

### Generation of Singlet Oxygen (^1^O_2_) (ABDA assay & SOSG)

The measurement of Co-CQDs capacity to generate singlet oxygen in different pH’s was monitored by measuring a decrease in fluorescence from the singlet oxygen sensitive and specific probe 6,10-anthracenediyl-bis(methylene) dimalonic acid (ABDA) when in the presence of Co-CQDs. Briefly, ABDA solution (100 µl, 40 µM) was introduced to solutions of Co-CQDs (0.12 – 0.0075%) at pH 5, 7 and 9. This solution was incubated at 37°C for 2 h and fluorescence was measured at λex / λem = 380 nm / 400–600 nm using a Tecan SPARK® spectrophotometer with black 66 welled plates. Singlet oxygen generation was determined from the decrease in fluorescence of the probe at 405 nm compared to time zero and as a fraction of the control (ABDA in PBS without nanoparticles at pH 5, 7 of 9).

To verify singlet oxygen production, we also conducted a singlet oxygen sensor green (SOSG, Thermo Fisher) assay. SOSG emits a strong green fluorescence upon selective oxidation by singlet oxygen. For the assay, 20 μl of SOSG solution in PBS (50 μM) was combined with varying Co-CQD concentrations 75-300 μg/ml in PBS and incubated at 37 °C for 4 hours. Endpoint fluorescence measurements were taken using a TecanSpark spectrofluorometer (λ_ex = 504 nm, λ_em = 530 nm. A control consisting of SOSG in PBS without nanoparticles was treated under identical conditions.

### Reactive Oxygen Species Generation Ascorbic Acid (AA) Oxidation

Ascorbic acid is susceptible to oxidation by ROS, producing dehydroascorbic acid. The ability of Co-CQDs to generate ROS was evaluated by monitoring the reduction in ascorbic acid absorbance at 266 nm in their presence. AA solutions (300 µM, 500 µl) prepared in PBS was combined with Co-CQDs dispersed in PBS (pH 7; 500 µl, 0.0025–0.02%) and incubated at 37°C for 2 hours. Absorbance spectra were recorded from 200 to 400 nm using a quartz cuvette. Ascorbic acid in PBS served as the negative control, while 15 mM H₂O₂ was used as a positive control.

### Hydrogen Peroxide Production - Amplex Red Assay

Hydrogen peroxide production by Co-CQD nanoparticles was evaluated using the Amplex Red assay according to the manufacturer’s protocol (Invitrogen, A22188). In brief, Co-CQDs (2.3–300 μg/ml) were incubated with 100 μM Amplex Red reagent and 0.2 U/ml horseradish peroxidase for 30 minutes at room temperature in the absence of light. A 15 mM H₂O₂ solution was used as a positive control, while ultrapure water served as the negative control. Fluorescence associated with H₂O₂ formation was recorded at λex/λem = 560 nm/580–650 nm.

### Membrane Polarization

MRSA cells (25 μl, OD_600_ = 0.10) were combined with 125 μl of 3,3′-dipropylthiadicarbocyanine iodide (diSC3-5) at a final concentration of 1.0 μM in HEPES buffer (10 mM) supplemented with glucose, KCl, and MgSO₄. The mixture was incubated for 30 minutes prior to the addition of Co-CQDs (37.5–1000 μg/ml). Fluorescence outputs were measured after 2 hours using a Tecan SPARK microplate reader, at λex/λem = 622 nm/674 nm.

## Microbiology

### Strains and Media

Bacterial strains used within this study were as follows:

*Staphylococcus aureus* subsp. aureus Rosenbach (methicillin and oxacillin resistant) – ATCC 43300, *Pseudomonas aeruginosa (Schroeter) Migula-PAO1 - BAA-47™*. Bacterial strains were routinely cultured in nutrient or Luria-Bertani broth and streaked onto nutrient or Luria-Bertani agar.

### Antimicrobial (Liquid culture)

Mid-exponential phase cultures of MRSA and PAO1 were exposed to varying concentrations of Co-CQDs (6–600 µg/ml) in nutrient broth and incubated at 37°C for 24 hours. Initial inoculum densities were 2.37 × 10⁶ CFU/ml for *P. aeruginosa* (PAO1) and 1.03 × 10⁶ CFU/ml for MRSA (*S. aureus)*.

To determine the minimum bactericidal concentration (MBC), 20 µl samples from each treatment were spread plated onto agar following the 24-hour incubation and cultured at 37°C. After 48 hours, colony-forming units (CFUs) were enumerated.

Metabolic activity was evaluated using a resazurin assay. Briefly, 30 µl of 0.015% (w/v) resazurin solution was added to 100 µl of each treated culture. After incubation for 3 hours, fluorescence was measured at λex/λem = 560 nm/685 nm.

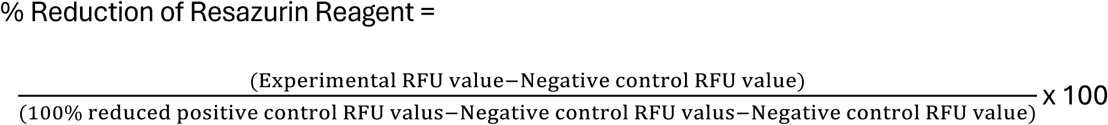

### Antimicrobial Live/Dead (BACLIGHT)

*Staphylococcus aureus* (2.37 × 10⁶ CFU) was applied to modified electrospun gels (prepared without the HPTS pH-sensitive probe) and allowed to infiltrate the matrix for 70 minutes. The gels were then submerged in LB broth and incubated at 37 °C for 16 hours. Following incubation, the hydrogels were removed, rinsed in PBS, and transferred to Ibidi 8-well plates.

Each gel was subsequently immersed in LB medium containing SYTO 9 and propidium iodide, prepared according to the manufacturer’s instructions (LIVE/DEAD BacLight Bacterial Viability Kit, Invitrogen). Bacterial cells were imaged using a Dragonfly 505 (Andor Technologies, Inc) confocal microscope with a 60 × oil immersion objective. Live cells were visualized at an excitation/emission of 480/500 nm, while dead cells were imaged at 490/635 nm.

### *In vivo* Wound Healing and Antimicrobial Assessment

All animal procedures were conducted with approval from the Norwegian Food and Animal Safety Authority (Mattilsynet; FOTS approval no. 30052). Surgical interventions were carried out under general anesthesia to reduce distress, and animals were randomly allocated to experimental groups.

Female Black C57BL/6 (8–10 weeks old) were prepared by shaving the dorsal surface, followed by chemical depilation using a commercial hair removal cream (Veet). Full-thickness excisional wounds were created using a 5 mm biopsy punch. Wounds were inoculated with MRSA (2.6 × 10⁶ CFU) and allowed to establish for up to 3 hours before treatment with 100 µl of Co-CQD containing hydrogel (∼12.5 µg/kg ±10%). Uninfected wounds irrigated with PBS served as the negative control, whereas untreated MRSA-infected wounds served as the positive control.

To limit contraction, a splint was adhered around the wound margins, and the site was covered with a transparent dressing (Tegaderm Film, 3M, 7000002875UPC). Each treatment group consisted of six animals. After 7 days, mice were euthanized and wound tissues were excised. Half of the samples (n=3) were homogenized in 1 ml sterile PBS, and 20 µl aliquots were spread plated on agar for bacterial enumeration by colony counting. The remaining samples were fixed, paraffin-embedded, and processed for hematoxylin and eosin (H&E) staining as well as immunolabeling analyses.

### Hematoxylin and Eosin (H&E) Staining

Wound tissue samples were fixed in 10% formalin, followed by dehydration through a graded ethanol series (70–100%). Specimens were then cleared in xylene and embedded in paraffin prior to sectioning. For H&E staining, sections were first dewaxed to remove paraffin and rehydrated. Slides were subsequently stained with hematoxylin, rinsed, counterstained with eosin, and rinsed again before undergoing dehydration and clearing. Finally, sections were mounted and imaged using an Olympus VS120 slide scanning system.

### Wound Tissue Immunofluorescence

Paraffin-embedded tissue sections (5 µm) were first deparaffinized in xylene (2 × 10 minutes) and subsequently rehydrated through a graded ethanol series (100% twice for 10 minutes each, 95% once for 10 minutes, and 70% twice for 10 minutes), followed by rinsing in distilled water (2 × 5 minutes). Antigen retrieval was carried out by heating samples in a pressure cooker at 98 °C for 12 minutes.

For MPO and iNOS staining, antigen retrieval was performed using a Tris-based unmasking solution (pH 9.0; Vector Laboratories, H-3301), while sodium citrate buffer (pH 6.0; Vector Laboratories, H-3300) was used for F4/80. After cooling to room temperature for 40 minutes, sections were blocked for 1 h with either 10% normal goat serum for F4/80 (Invitrogen, 50197Z) or 5% bovine serum albumin in PBS for MPO and iNOS.

Primary antibodies were applied overnight at 4 °C, with F4/80 (Cell Signaling, 70076) and MPO (R&D Systems, AF3667) diluted 1:200, and iNOS (Abcam, ab15323) diluted 1:100. Following washes in PBS containing 0.05% Tween-20, sections were incubated in the dark for 1 hour with fluorescent secondary antibodies (1:200; Alexa Fluor 647, Invitrogen A21244; Alexa Fluor 546, Invitrogen A11056). After final washing steps, slides were mounted using ProLong Diamond antifade reagent with DAPI (Invitrogen, P36970). Imaging was performed using an Olympus VS120 slide scanner with a 40× objective.

### Trichrome Staining

Collagen-rich connective tissue was visualized using the Trichrome Stain kit (Abcam, ab150686). Briefly, 5 mm tissue sections were deparaffinized and rehydrated following the protocol outlined in the tissue immunofluorescence methods. Sections were then stained in accordance with the manufacturer’s guidelines. After staining, slides were cleared in xylene and mounted using Eukitt quick-hardening mounting medium (Sigma-Aldrich, 03989). Imaging was performed with an Olympus VS120 slide scanner at 40× magnification.

### Statistics and Reproducibility

All microscopy experiments, including transmission electron microscopy (TEM) and confocal imaging, were performed in three independent replicates. Error bars indicate the standard deviation of the mean. Mice were randomly allocated to treatment groups and further randomized for either wound bacterial load assessment or histological analysis. Statistical comparisons among groups were conducted using one-way or two-way analysis of variance (ANOVA), followed by Tukey’s post hoc test. In specific assays, an unpaired t-test with Welch’s correction was applied. All analyses were performed using a 95% confidence level, with statistical significance defined as p < 0.05.

## Acknowledgments

We would like to acknowledge (1) The Molecular Imaging Centre (MIC), Department of Biomedicine, University of Bergen (Norway) for use of their electron and confocal microscopes. (2) The facilities, and the scientific and technical assistance of the RMIT University’s Microscopy & Microanalysis Facility, a linked laboratory of the Microscopy Australia, enabled by NCRIS. (3) The facilities, and the scientific and technical assistance of Microscopy Australia (https://ror.org/042mm0k03) and the Australian National Fabrication Facility (ANFF) (https://ror.org/04ypnrn45), enabled by NCRIS and the government of South Australia at Flinders Microscopy and Microanalysis (https://ror.org/04z91ja70), Flinders University (https://ror.org/01kpzv902).

---

**Supplementary figure 1:**
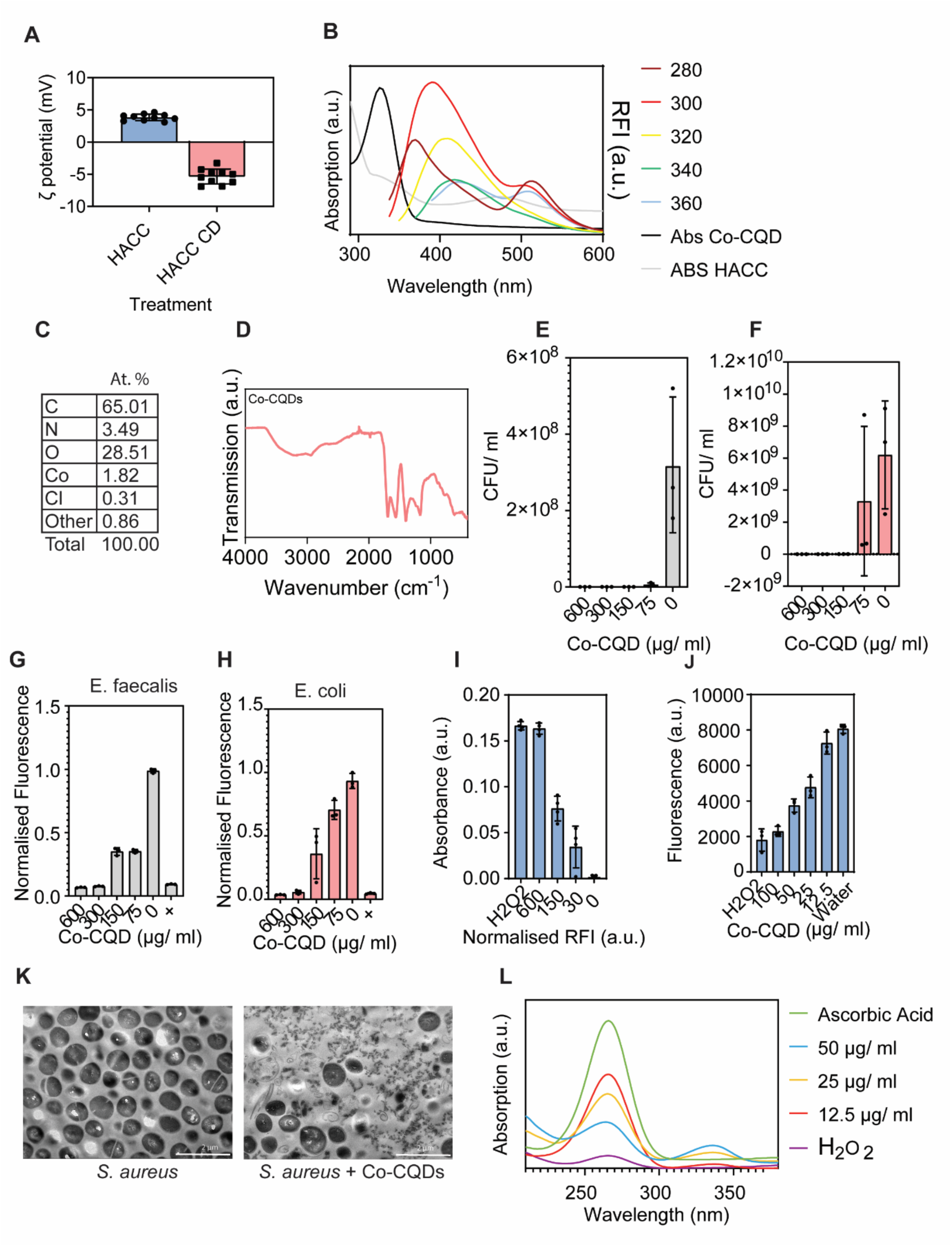
**Co-CQD nanoparticle characterisation and antimicrobial activity.**(A) Co-CQD particle and hexamine cobalt chloride precursor charge. (B) Absorbance and fluorescence characteristics of Co-CQD nanoparticles. (C) XPS elemental composition (Wt. %) of Co-CQDs. (D) FTIR spectra of Co-CQDs. (E) Bactericidal activity of Co-CQDs against *Enterococcus faecalis* bacteria. (F) Bactericidal activity of Co-CQDs against *Escherichia coli* bacteria. (G) Metabolic activity of *Enterococcus faecalis* bacteria exposed to Co-CQDs. (H) Metabolic activity of *Escherichia coli* bacteria exposed to Co-CQDs. (I) Hyperpolarisation of MRSA with varying Co-CQD concentrations. (J) Extracellular alkaline phosphatase in growth medium with MRSA cells exposed to varying Co-CQD concentrations. (K) Transmission electron microscope images of MRSA cells without and with exposure to Co-CQDs. (L) Ascorbic acid oxidation by Co-CQDs between 12.5 – 50 µg/ml concentration.

**Supplementary figure 2:**
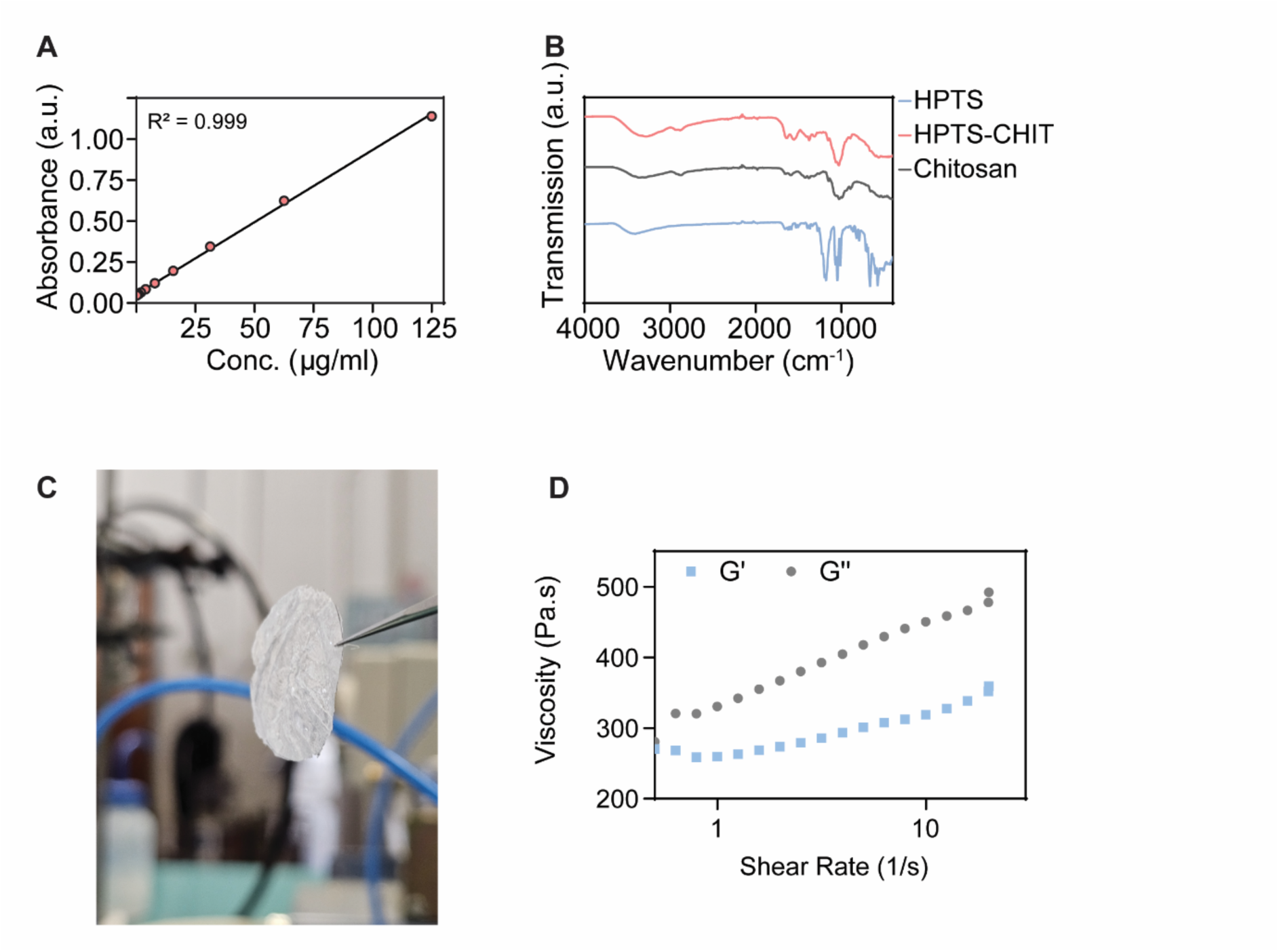
Fluorescent HPTS/ chitosan nanoparticle and hydrogel characterisation. (A) Linear region of HPTS/ chitosan nanoparticle absorbance with varying concentrations of the probe. (B) FTIR spectra of HPTS/chitosan nanoparticles along with the precursors used to synthesize them (HPTS dye and chitosan (C) Freeze dried hydrogel thin film. (D) Rheological frequency sweep of the hydrogel. +/-Gel refers to the hydrogel containing Co-CQD nanoparticles (+) and hydrogel without Co-CQD nanoparticles (-) respectively.

**Supplementary figure 3:**
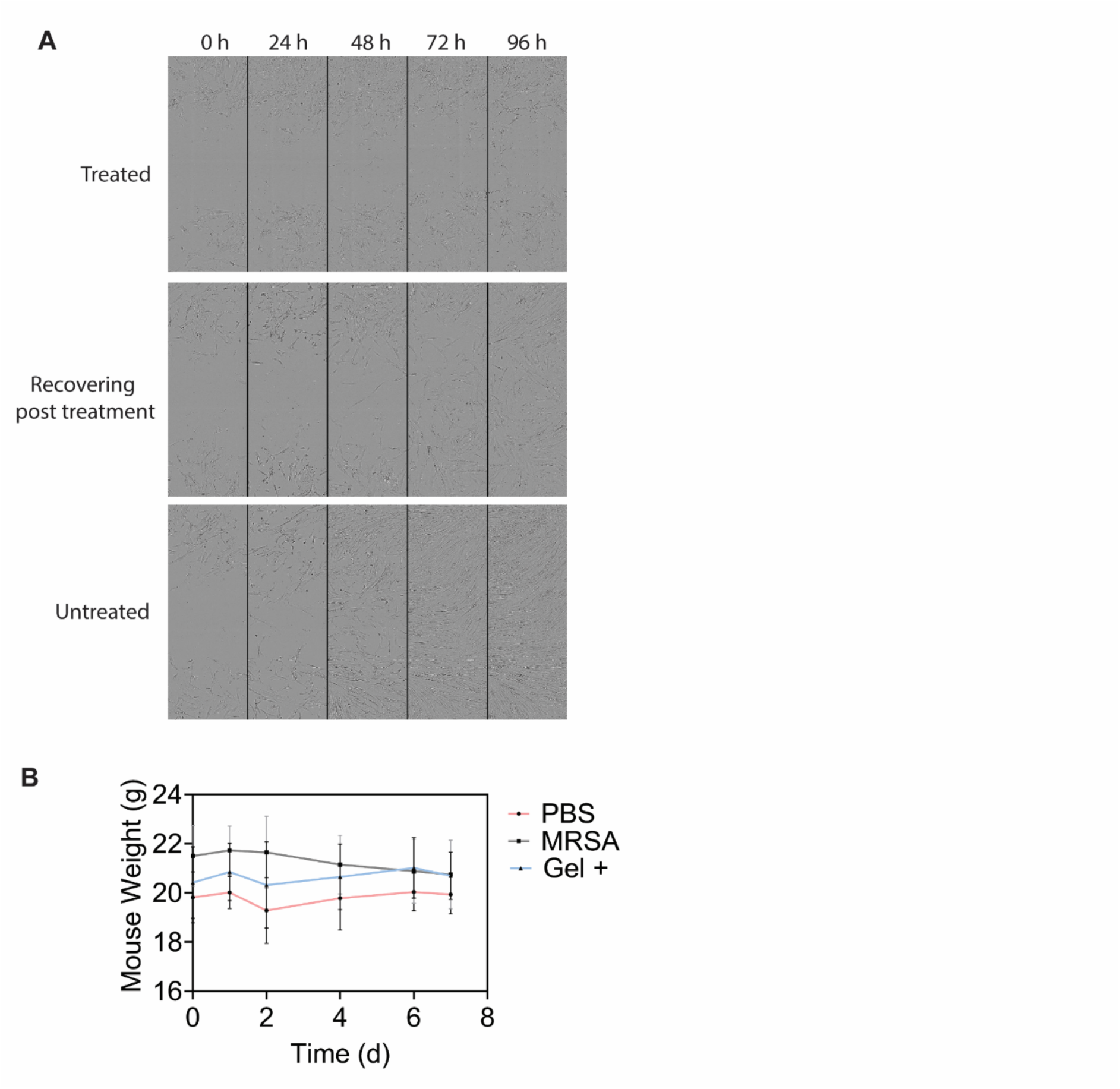
Biocompatibility of hydrogel. (A) Scratch wound assays of primary dermal fibroblast cell growth following being treated with Co-CQDs (500 µg/ ml), cell media replaced after being exposed to Co-CQDs (500 µg/ ml) for 24 h, and untreated cells over 66 h.

